# Skin microbiota variation in Indian families

**DOI:** 10.1101/2023.12.09.570904

**Authors:** Renuka Potbhare, Ameeta Ravikumar, Eveliina Munukka, Richa Ashma, Leo Lahti

## Abstract

**Background:** In Indian culture, extended families have symbolized our tradition. Families often encompass members spanning multiple generations cohabiting the same household, thereby sharing ethnicity, genetics, dietary habits, lifestyles, and living conditions. The joint or extended family setup provides an opportunity to compare variations in microbiota composition within and between families. While previous research has demonstrated that skin microbiota can be influenced by factors such as ethnicity, geography, diet, age, and sex, its associations among Indian family members that may share also genetic background remains largely unexplored.

**Methods:** The present study involved seventy-two individuals from fifteen families in two geographical regions of Maharashtra, India. Bacterial DNA was extracted from axillary sweat samples, followed by sequencing of V3-V4 regions of the 16S rRNA. The generated taxonomic profiles were used to quantify microbiota diversity and similarities in skin microbiota composition within and between families, taking into account factors such as genetic relatedness, diet, sex, age, geographical location, and co-habitation.

**Results:** The skin microbiota composition typically comprised Firmicutes, Proteobacteria, and Actinobacteria phyla. Notably, the Shannon alpha diversity was moderately associated with dietary habits and geographical location (Kruskal-Wallis; FDR<0.1), whereas no significant differences were observed for other key factors such as age, location, or sex. A significant association was also observed between taxonomic composition and shared familial membership (p=0.001; PERMANOVA), with a borderline significant association with geographical location (p=0.07). When within and between family comparisons were investigated across three generation (G1-G2, G2-G3 and G1-G3), no significant differences were observed, however, in general skin microbiota was more similar within than between families.

**Conclusion:** This study underscores the diversity and commonalities in skin microbiota composition within and between families. We observed that every family has a unique skin microbiota and among the various covariates, significant association was observed for diet and geographical location. Our study highlights that family relations may have specific associations with skin microbiota composition and diversity. Further studies with larger sample sizes will help to elucidate the relative contributions of shared co-habitation and genetic backgrounds.

## Introduction

Skin is an epithelial interface mediating the interaction between the internal and external body environment and it provides a first line of defence against toxins and invasion of various pathogens (Gallo, 2017; Coates et al., 2019; & Williams et al., 1998). The human skin also offers a niche for diverse communities of bacteria, fungi, viruses, and other micro-organisms collectively known as skin microbiota (Byrd et al., 2018). These microbes are classified as resident or transient species based on their survival on the human skin surface (Grice et al., 2008). Resident species live longer on the skin, are not harmful, and mainly belong to *Propionibacterium*, *Corynebacterium*, *and Staphylococcus* (Scholz & Kilian, 2016). However, these bacteria can play a beneficial or opportunistic role depending on the skin microenvironment (Findley and Grice, 2014). Whereas transient species like *Bacillus species*, *Staphylococcus aureus*, and *Pseudomonas aeruginosa* are short-term residents and less stable on the skin as it readily influenced by changes in the physio-chemical properties of the skin (Bojar & Holland, 2002).

Early-life microbial communities are typically colonized by maternal microbiota (Yao et al., 2021, Dominguez-Bello et al., 2010); later dynamic shifts are observed in the microbiota, and microbiota maturates gradually during the first years of life. Microbiota composition is notably diverse on healthy skin and affected by multiple factors like age (Chaudhari et al., 2020; Kim et al., 2019), diet (Salem et al., 2018), geography (Ross et al., 2018), sex (Byrd et al., 2018, Li et al., 2019), and environment (Peng et al., 2020, Lehtimäki et al., 2017). Skin type (dry, moist, sebaceous), skin site and skin parameters such as pH, sebum levels, number of hair follicles and gland also contributes in shaping skin microbiota communities (Oh et al., 2014, Cho and Eom, 2021). Despite these endogenous and exogenous factors, microbiota varies between individuals and can be influenced by genetics (Si et al., 2015, Skowron et al., 2021). Additionally, the host lifestyle, hygiene (Riverain-Gillet et al., 2020), and use of cosmetics (Wallen-Russell., 2018, Salverda et al., 2013) influence the abundance and composition of microbial communities inhabit skin. In addition to these factors skin microbiome varies based on phenotypic differences between host and external factors such as environmental exposure (Ruuskanen et al., 2022).

However, understanding the role of genetic relations remains unclear and has not been interrogated, and skin microbiota studies have largely focused on associations such as cohabitation (Ross et al., 2017), human-pet association (Song et al., 2013), captivity, and indoor environment etc (Lax et al., 2014). To date, research about the skin microbiota associations among genetically related individuals has been limited.

Various animal models and human samples have been used in previous studies stating significant association of host genetic effects with gut microbiota composition (Nichols & Davenport., 2021, Zhao et al., 2013; Goodrich et al., 2014, Turpin et al., 2016). Also, the gut microbiota studies reported the passing of some gut microbiota to the next generation maintaining heritability pattern, and the underlying role of genetic components in mutualistic host-microbiota interactions (Turpin et al., 2016, Hughes et al., 2020, Sanna et al., 2022). However, the relationship between an individual’s genetics and skin microbiota composition and diversity is poorly understood.

There is evidence supporting vertical transmission of skin microbiota from mother to child which depends on method of childbirth, maternal health, breastfeeding, close contact and other factors, which play a vital role in shaping skin microbiota of an infant (LaTuga et al., 2014, Rapin et al., 2023). Dominguez-Bello et al., 2010 showed that babies born vaginally exhibit a microbiota composition in their birth canal that is similar to that of their mothers while infants delivered by cesarean-section acquire bacterial communities resembling skin surfaces (Dominguez-Bello MG et al., 2010). Furthermore, heredity is a significant factor in the transmission of microbiota from one generation to the next. A recent study demonstrated that individuals who are genetically related exhibit a higher degree of similarity in their gut microbiota compared to those who are not genetically related (Grieneisen et al., 2021). However, it is important to consider the effect of host environment on the degree of heritability (Grieneisen et al., 2021). Therefore, it is crucial to consider factors such as host genetics, interactions between hosts, and the shared living environment of host.

Joint or extended families have been a feature of Indian society since ancient times (Karve, 1961; Mullaiti et al., 1995). This comprises a group of multi-generational people who are related to each other, living together under one roof, preparing food at a common hearth, and who share similar dietary habits, hygiene, and house environmental conditions (Karve, 1953; Chadda et al., 2013). Thus, the study of skin microbiota similarity between family members provides an opportunity to understand skin microbiota variation associated with the genetic relatedness of individuals within the same family but this has to be carefully controlled for the shared effects of diet, co-habitation, and environmental conditions.

We investigated differences in skin microbiota taxonomic diversity and composition between genetically related and unrelated individuals within and between families across three generations considering various background factors. We analyzed 72 healthy individuals from fifteen families wherein three generations live together under the same roof.

## Materials and methods

### Study design and subject enrolment

Seventy-two volunteers comprising fifteen families were enrolled in the present study from two districts of Maharashtra, India, viz, Pune (altitude: 1,840 ft; longitude:73°51ʹ19.26ʺE; latitude:18°31ʹ10.45ʺN) and Ahmednagar (altitude:2129 ft; longitude: 74°44ʹ58.53ʺE; latitude:19°5ʹ42.75ʺN). Members from the same family have similar dietary habits, vegetarian or mixed, except two families (**Table 1**). We classified selected volunteers of each family into three generations by age, generation-1 (G1, age 65-91 yrs.), generation-2 (G2, 41-63 yrs.), and generation-3 (G3, 13-30 yrs.). The volunteers fulfilled the following conditions and criteria: a) Each family must comprise individuals from three different generations (at least one member per generation), i.e., grandmother or grandfather (age 65-91 yrs.), mother or father (age ranged within 41-63 yrs.), daughter or son respectively (age 13-30 yrs.). b) All members of the same family must share a household. c) All volunteers were self-identified as healthy. The volunteers were instructed to avoid using deodorants, skin ointments, and soaps before 12-15 hrs of sampling, shaving of axilla at least two days before sampling, tobacco, smoking, and alcohol consumption, and certain food items like onion, garlic, chilies. The demographics of the fifteen enrolled families are described in **Figure 1** and **Table 1**.

**Figure 1:**
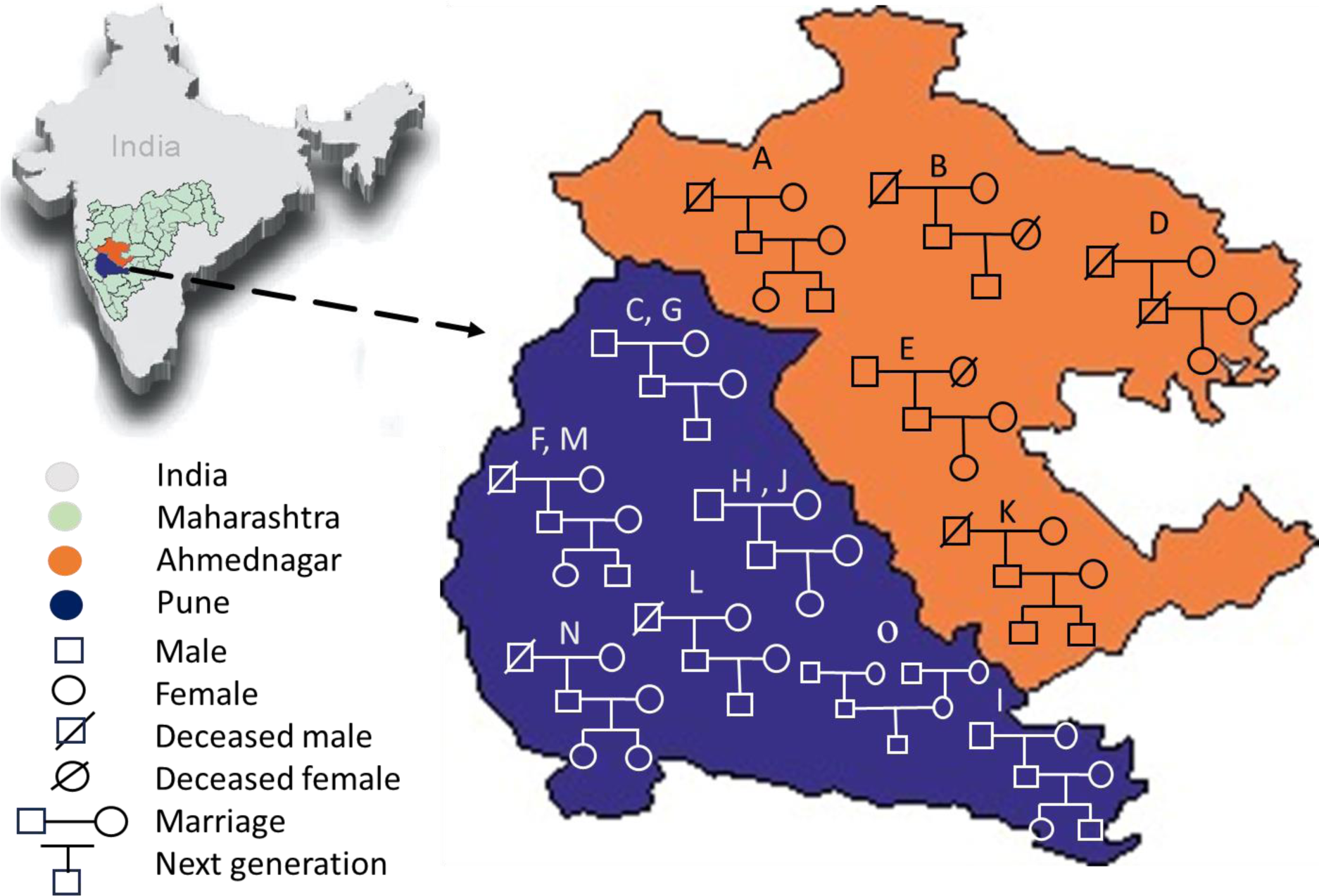
Demographics of the enrolled families (A-O) represents geography-wise family distribution and pedigree structure across three generations.

**Table 1:** Information on the enrolled families.

| Family | Members | Generation |  |  | Diet |  | Sex |  | Location |  | Age |  |  |
| --- | --- | --- | --- | --- | --- | --- | --- | --- | --- | --- | --- | --- | --- |
|  |  | G1 | G2 | G3 | Vegetarian | Mixed | Male | Female | Pune | Ahmednagar | Adult | Middle age | Elderly |
| (n=15) | (n=72) | (n=32) | (n=28) | (n=21) | (n=57) | (n=15) | (n=33) | (n=39) | (n=52) | (n=20) | (n=25) | (n=26) | (n=21) |
| A | 5 | 1 | 2 | 2 | 5 | 0 | 2 | 3 | 0 | 5 | 1 | 2 | 2 |
| B | 3 | 1 | 1 | 1 | 3 | 0 | 2 | 1 | 0 | 3 | 1 | 1 | 1 |
| C | 5 | 2 | 2 | 1 | 5 | 0 | 3 | 2 | 5 | 0 | 2 | 2 | 1 |
| D | 3 | 1 | 1 | 1 | 3 | 0 | 0 | 3 | 0 | 3 | 1 | 1 | 1 |
| E | 4 | 1 | 2 | 1 | 0 | 4 | 2 | 2 | 0 | 4 | 2 | 1 | 1 |
| F | 5 | 1 | 2 | 2 | 1 | 4 | 2 | 3 | 5 | 0 | 1 | 2 | 2 |
| G | 5 | 2 | 2 | 1 | 5 | 0 | 3 | 2 | 5 | 0 | 2 | 2 | 1 |
| H | 5 | 2 | 2 | 1 | 5 | 0 | 2 | 3 | 5 | 0 | 2 | 2 | 1 |
| I | 6 | 2 | 2 | 2 | 6 | 0 | 3 | 3 | 6 | 0 | 2 | 2 | 2 |
| J | 5 | 2 | 2 | 1 | 0 | 5 | 2 | 3 | 5 | 0 | 2 | 2 | 1 |
| K | 5 | 1 | 2 | 2 | 5 | 0 | 3 | 2 | 0 | 5 | 2 | 1 | 2 |
| L | 4 | 1 | 2 | 1 | 4 | 0 | 2 | 2 | 4 | 0 | 1 | 2 | 1 |
| M | 5 | 1 | 2 | 2 | 5 | 0 | 2 | 3 | 5 | 0 | 1 | 2 | 2 |
| N | 5 | 1 | 2 | 2 | 3 | 2 | 1 | 4 | 5 | 0 | 1 | 2 | 2 |
| O | 7 | 4 | 2 | 1 | 7 | 0 | 4 | 3 | 7 | 0 | 4 | 2 | 1 |

Volunteers were informed about the study goals and provided a questionnaire to obtain information about lifestyle and medication history. The questionnaire covered the history of dermatological disease, use of antibiotics, long-term medication history, diet habits, amount of physical exercise, etc. The experiments involved in the study were approved by an appropriate institutional human ethics committee of Savitribai Phule Pune University (Letter No. EC/2016/15, Dated: 1^st^ October 2016).

### Sample collection and microbial DNA extraction

Skin sweat swabs were collected from the axillary region using sterile cotton swabs moistened with PBS solution (1X). Cotton buds were scrubbed multiple times in upward and downward directions to capture the axillary microbiota. Collected sweat samples were stored at −20°C/−80°C until genomic DNA extraction (2-3 months) to prevent degradation of samples. Bacterial DNA was extracted from each sample using methods described previously (Potbhare et al., 2022). In brief, the bacterial cells were lysed using lysis buffer (0.5M EDTA, 0.5 M Tris-HCl, 0.1v/v Triton X, 8% sucrose), followed by double purification with Phenol: chloroform: iso amyl alcohol (25:24:1) method described in (Sambrook, Fritsch & Maniatis, 1989). The mixture was precipitated by washing several times with 100% and 70% ethanol. The resulting nucleic acid mixture was confirmed for DNA extraction by loading on gel electrophoresis technique and assessed for the quality and quantity using a nanodrop spectrophotometer at 260 nm and 280 nm absorbance. Further, these DNA samples were subjected to *16S rRNA* sequencing of V3 and V4 regions on the Illumina MiSeq platform.

### Library preparation and *16S rRNA* gene sequencing

Universal prokaryotic primers of V3-V4 regions were used to amplify the *16S rRNA* gene. Polymerase Chain Reactions (PCR) were performed using KAPA HiFi HotStart Ready Mix® (KAPA biosystems, Boston, USA) with the following thermal parameters: initial denaturation at 95°C for 3 min, followed by 25 cycles of 95°C for the 30 sec; 55°C for the 30 sec; and 72°C for 30 sec. The resulting amplicons were cleaned and purified with AMPure XP beads using PureLink^TM^ PCR purification kit® (Invitrogen, Waltham, MA, USA). Further, dual indices and adaptors were linked to the amplicon using the Nextera XT Index Kit® (Illumina, San Diego, CA, USA) following the thermal cycler program: initial denaturation at 95°C for 3 min, followed by eight cycles of 95°C for the 30 sec; 55°C for the 30 sec; and 72°C for 5 min. These adaptor-ligated libraries were purified with AMPure XP beads using PureLink^TM^ PCR purification kit® (Invitrogen, Waltham, MA, USA). The quantitation of PCR products was measured using Qubit^TM^ dsDNA HS Assay Kit® (Invitrogen, Waltham, MA, USA) on a 2.0 fluorometer (Life Technologies, Waltham, MA, USA). These products were pooled into equal molar proportions and sequenced on Illumina MiSeq V2 standard flow cell (Illumina, San Diego, CA, USA) for 2*300 bp pair-end chemistry according to the manufacturer’s instructions. A 5% PhiX control (Illumina, San Diego, CA, USA) along with positive (DNA sample extracted from the healthy gut) and negative controls (MilliQ water sample) were included in the final run to rule out the possibility of contamination from the experimental materials.

### Bioinformatics

The bacterial *16S rRNA* gene sequences in FASTQ format were aligned using Illumina BaseSpace toolkit. The primer and adapter sequences were trimmed from the raw sequences. The sequences were filtered after trimming the 3′ end with the quality score (Q) 30. This eliminated PCR-generated chimeras, contaminants, and low-quality reads from the sequences. In order to generate the OTU table, RefSeq RDP 16S v3 May 2018 DADA2 was used for taxonomic identification (Callahan et al., 2016). Altogether 5 710 132 reads, ranging between 30 281-134 873 reads with an average ∼80 000 reads per sample were generated. The amplicon sequence reads were then grouped into 1,070 OTUs based on <98% sequence similarity cutoff. Then the data was imported into *TreeSummarizedExperiment* data container in R in order to organize the taxonomic profiling data and associated metadata on samples and features (v. 2.6.0) (Huang et al., 2021).

### Statistical analysis

We quantified alpha diversity with Shannon index, and associated this with multiple covariates including diet, age, sex, geographical location, and family. Further, dissimilarities in taxonomic composition between individuals (beta diversity) was calculated with Permutational Multivariate Analysis of Variance (PERMANOVA) with 999 permutations using Bray-Curtis dissimilarity (Anderson, 2001). Principal coordinates analysis (PCoA) based on Bray-Curtis dissimilarity was used to visualize the distribution of taxonomic composition among the study population. The beta diversity measurements were calculated using the *vegan* R package (Oksanen et al., 2020). In order to investigate the similarity in the microbiota composition among family members across three generations, we compared individuals from G1-G2 (grandparent-parent), G2-G3 (parent-children), and G1-G3 (grandparent-children) and calculated within and between family (dis)similarities using Bray-Curtis index. For this analysis, we selected one member from each generation of the same family and calculated pair-wise within family similarity. For between family differences, we selected member of each generation from different families and calculated pair-wise between family (dis)similarity. Similarly, familywise inter-generational analysis was performed by selecting two members per generation (G1-G2 or G2-G3) using Bray-Curtis index. We further extended G2-G3 (parent-child) analysis, and separately compared father-child and mother-child skin microbiota similarity so as to obtain baseline (dis)similarity. The p-values were adjusted for multiple testing with Benjamini-Hochberg False Discovery Rate (FDR) correction for all the analysis. Comparisons with FDR<0.05 were considered significant, and with FDR<0.1 moderately significant. We limited all comparisons within same city, age, and sex in order to control the potential association of these factors with skin microbiota composition.

The analyses were carried out at genus level unless otherwise mentioned. All analyses and visualisations were carried out in R (version 4.2.2). The primary R packages were *mia* (1.6.0) and *miaViz* (1.6.0) (Ernst et al., 2022).

## Results

### Alpha diversity

We compared geographical location, diet, age, and sex with respect to Shannon alpha diversity. The diet and geographical location are moderately associated with alpha diversity (Kruskal-Wallis test, FDR <0.1). Differences with the other factors were not significant (**Figure 2**) (age: p=0.73; sex p=0.73; Kruskal-Wallis test).

**Figure 2:**
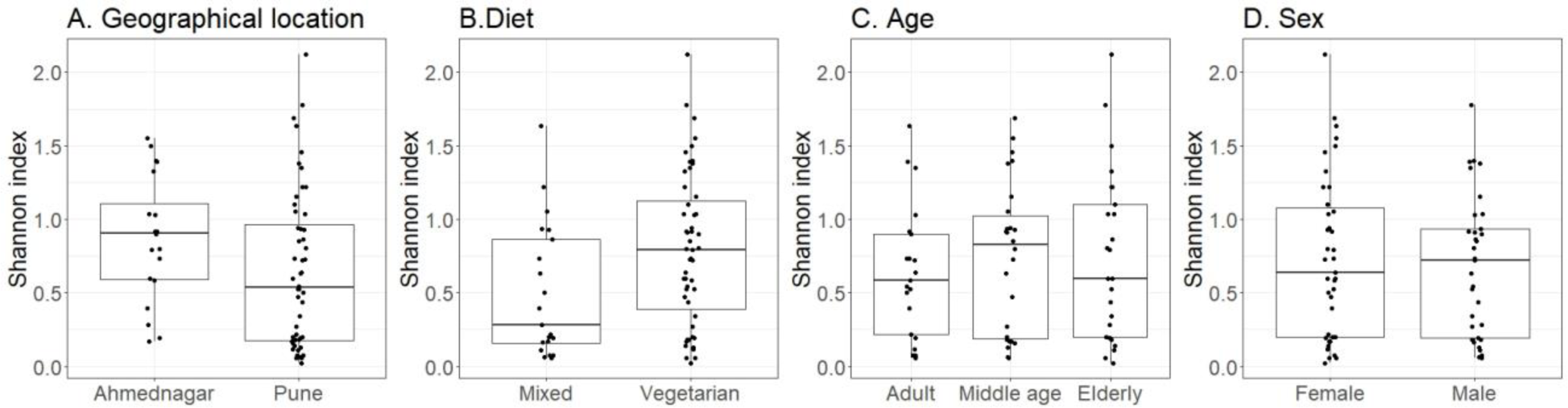
Alpha diversity (Shannon index) variation of the skin microbiota and its associations with A) location (Kruskal-Wallis test, p=0.08); B) diet (p=0.08); C) age (p=0.73); D) sex(p=0.73).

### Beta diversity

We observed significant association of skin microbiota with family (p=0.001) and borderline significant association with geographical location (p=0.07). We did not observe significant association between taxonomic composition and diet (p=0.93), age (p=0.36), or sex (p=0.61) (PERMANOVA; **Table 2**). Sample similarity is illustrated with principal coordinates analysis in **Figure 3**. Further, distance-based redundancy analyses (dbRDA) performed on major factors highlighted significant difference with geographical location (explained variance 18.4%; p=0.01), but not with diet (0.8%, p=0.5), age (1.8%, p=0.6) or sex (0.9%, p=0.5), as illustrated in **Figure 4**.

**Figure 3:**
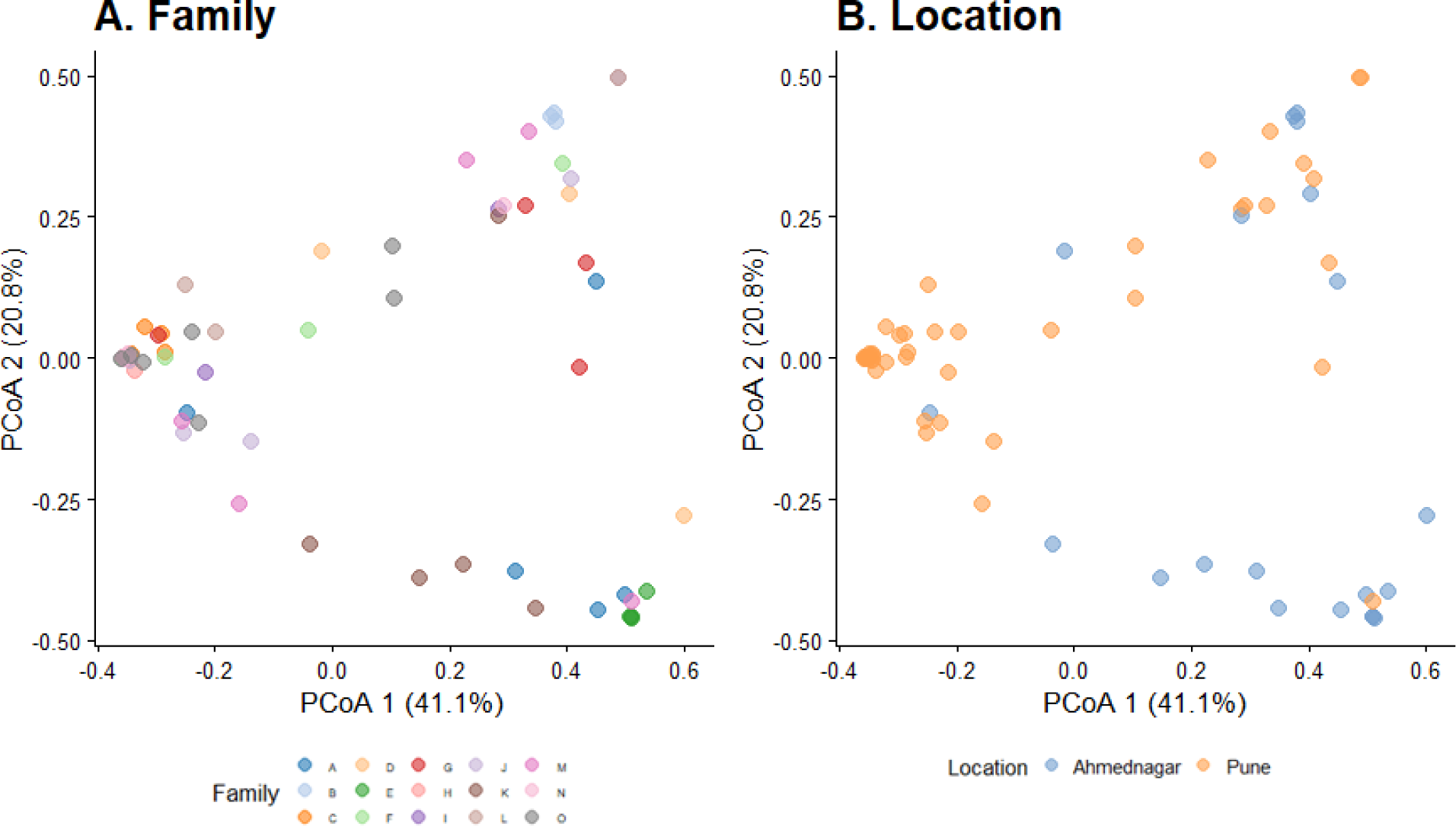
Dissimilarity of skin microbiota composition illustrated on Principal Coordinates Analysis ordination (PCoA; Bray-Curtis). The color indicates A) family; B) location.

**Figure 4:**
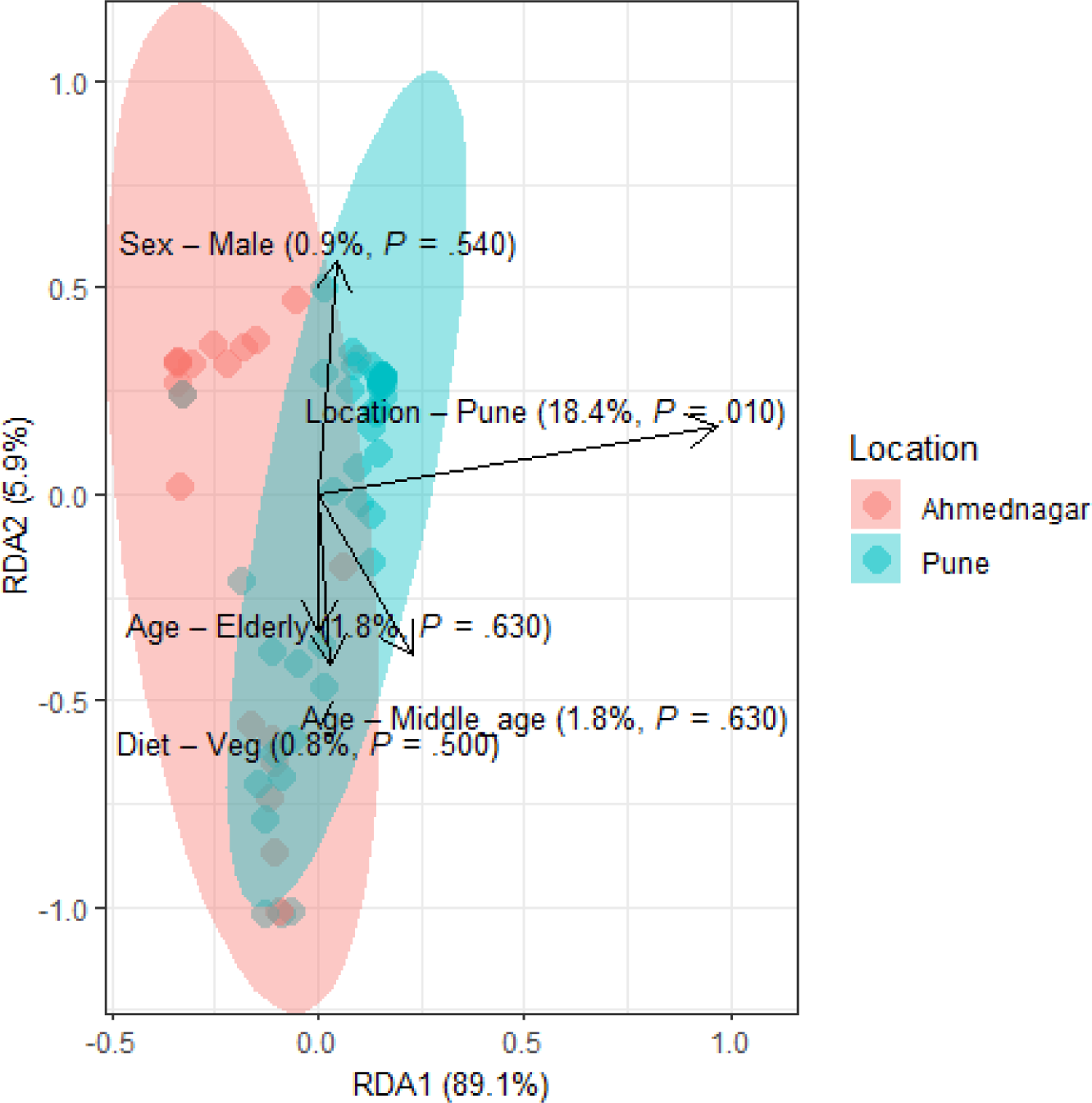
Distance-based redundancy (dbRDA) analysis of skin microbiota with major confounding factors (Bray-Curtis dissimilarity), location (18.4%; p=0.01**), diet (0.8%, p=0.5), age (1.8%, p=0.6) and sex(0.9%, p=0.5)

**Table 2:** Association between the skin microbiome taxonomic composition with key covariates (PERMANOVA; Bray-Curtis dissimilarity; Benjamini-Hochberg adjustment for multiple testing).

| Factor | R2 | p-value |
| --- | --- | --- |
| Diet | 0.00 | 0.93 |
| Age | 0.02 | 0.36 |
| Sex | 0.00 | 0.61 |
| Location | 0.19 | 0.07 |
| Family | 0.29 | 0.00 *** |

Note- The p-value indicates level of significance $<0.001$ \*\*\*, $<0.01$ \*\*, $<0.05$ \*, 0.1 ns

### Taxonomy profiling of skin microbiota

Skin microbiota observed in our study population comprised a total of 36 phyla, 84 classes, 191 orders, 326 families and 1071 genera. Most of these observed groups were rare and observed only in very few sample at a low abundance. Of the 36 phyla observed across the families, there were three most dominant phyla in the studied population viz., *Firmicutes* (prevalence 100%, mean relative abundance 73%), *Proteobacteria* (97.2%, 23.8%), and *Actinobacteria* (90.2%, 3.2%) having >1% prevalence and a detection threshold of 0.1% (**Table 3**). We further filtered the observed 1071 genera by setting 0.1% detection threshold and >1% prevalence, which resulted 73 genera. The remaining genera were grouped as ‘other’. Out of 73 genera the most dominant six genera with highest mean abundance were *Staphylococcus* (prevalence 100%, mean relative abundance 50.9%), *Bacillus* (87.5%, 16.3%), *Pseudomonas* (61.1%, 9.2%)*, Anaerococcus* (38.9%, 1.7%), *Corynebacterium* (73.6%, 1%), and *Dermabacter* (40.2%, 0.4%) (**Table 4**).

**Table 3:** The most prevalent phyla on skin microbiome with the mean relative abundance, median, and prevalence (detection threshold=0.1%, prevalence>1%).

| Phylum | Mean<br>relative<br>abundance<br>(%) | Median<br>(%) | Prevalence<br>(%) |
| --- | --- | --- | --- |
| *Firmicutes* | 73.0 | 93.5 | 100.0 |
| *Proteobacteria* | 23.8 | 2.6 | 97.2 |
| *Actinobacteria* | 3.2 | 1.2 | 90.2 |

**Table 4:** The most prevalent genera on skin microbiome with the mean relative abundance, median, and prevalence (detection threshold=0.1%, prevalence>1%).

| Genus | Mean<br>relative<br>abundance<br>(%) | Median<br>(%) | Prevalence<br>(%) |
| --- | --- | --- | --- |
| *Staphylococcus* | 50.9 | 60.2 | 100.0 |
| *Bacillus* | 16.3 | 0.3 | 87.5 |
| *Pseudomonas* | 9.2 | 0.1 | 61.1 |
| *Anaerococcus* | 1.7 | 0.0 | 38.9 |
| *Corynebacterium* | 1.0 | 0.2 | 73.6 |
| *Dermabacter* | 0.4 | 0.0 | 40.2 |

The geographical location-wise distribution of the three most prevalent phyla represented in heatmap (**Figure 5A**). We further examined location-wise taxonomic composition at the genus level for families. This revealed particularly high abundances of *Staphylococcus* among all families, and *Bacillus* as the most dominant genus in Ahmednagar. Overall, the abundances of these dominant genera varied across families and geographical locations (**Figure 5 b-c**).

**Figure 5:**
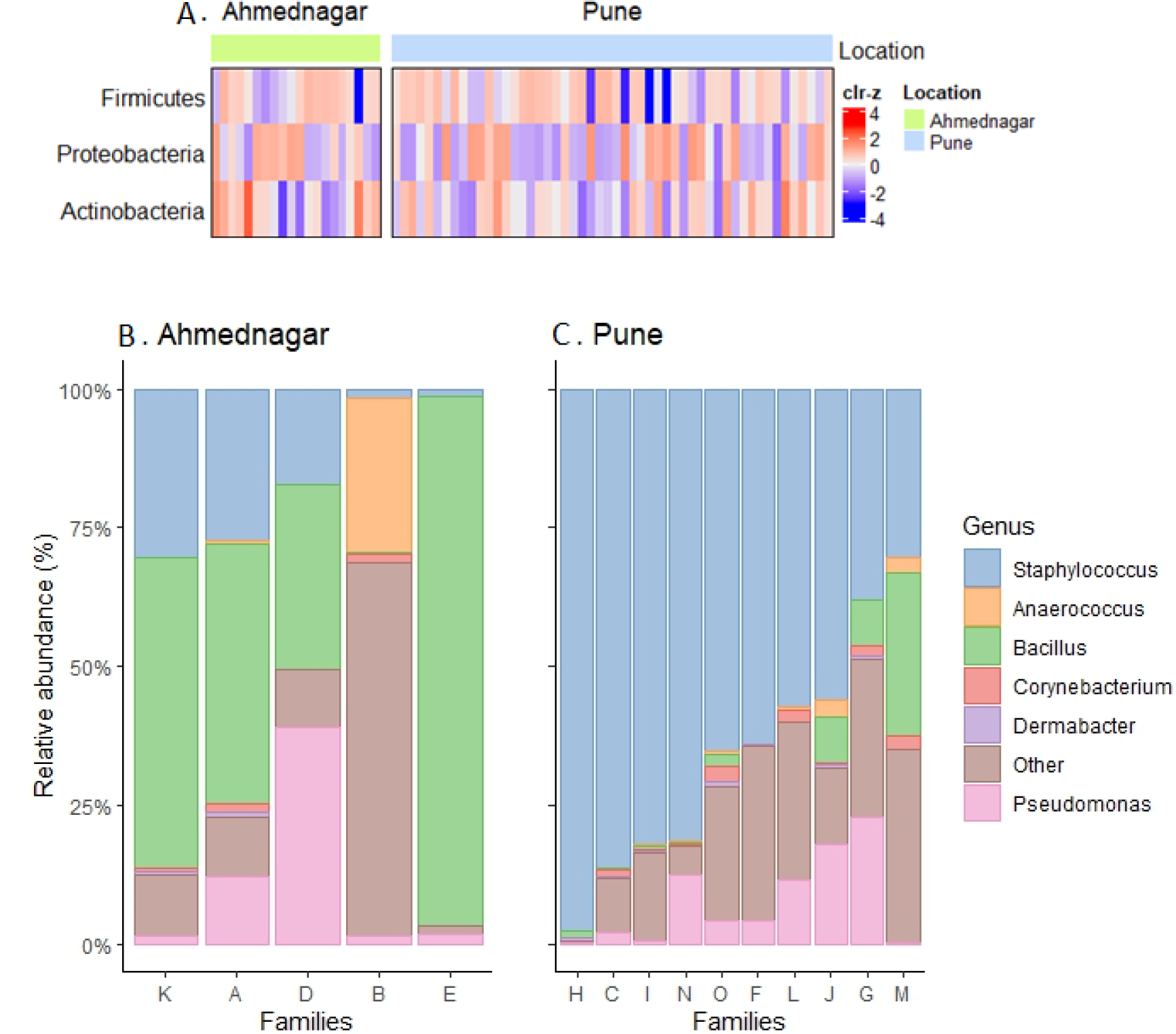
A) Heatmap representing the abundance variation for the most prevalent phyla on skin microbiota for the two locations (detection threshold=0.1%, prevalence>1%; data transformed with clr for samples, followed by z transformation on the taxonomic features); Location-wise relative abundances of the most prevalent genera in families. B) Ahmednagar (n = 5), C) Pune (n = 10) The “Other” represents the sum of relative abundances for the less prevalent genera. The samples are ordered based on Staphylococcus abundance.

### Inter-generational analysis of skin microbiota across families

Our analysis did not reveal significant differences in the microbiota composition between generations: G1-G2 (grandparent-parent), G2-G3 (parent-children), and G1-G3 (grandparent-children) for both within and between family comparisons (Kruskal-Wallis test, FDR<0.1, **Figure 6A-C**). Furthermore, when we conducted a separate analysis of G2-G3 (parent-child) relationship, we again did not find significant differences within and between families for father and mother, (Kruskal-Wallis test, FDR<0.1, **Figure 6 D-E**). On average children exhibited a greater degree of microbiota similarity with their parents than parents from other families. Also, we did not observe significant inter-generational differences within family (supplementary figure 1). We also investigated whether children shared a higher taxonomic similarity with their father compared to their mother but the difference was not significant (Kruskal-Wallis test, FDR<0.1, **Figure 6 F**). This parent-child analysis was conducted on the thirteen families that included both parents and children regardless of their sex. We excluded two families from Ahmednagar due to death of one of the parents in generation 2.

**Figure 6:**
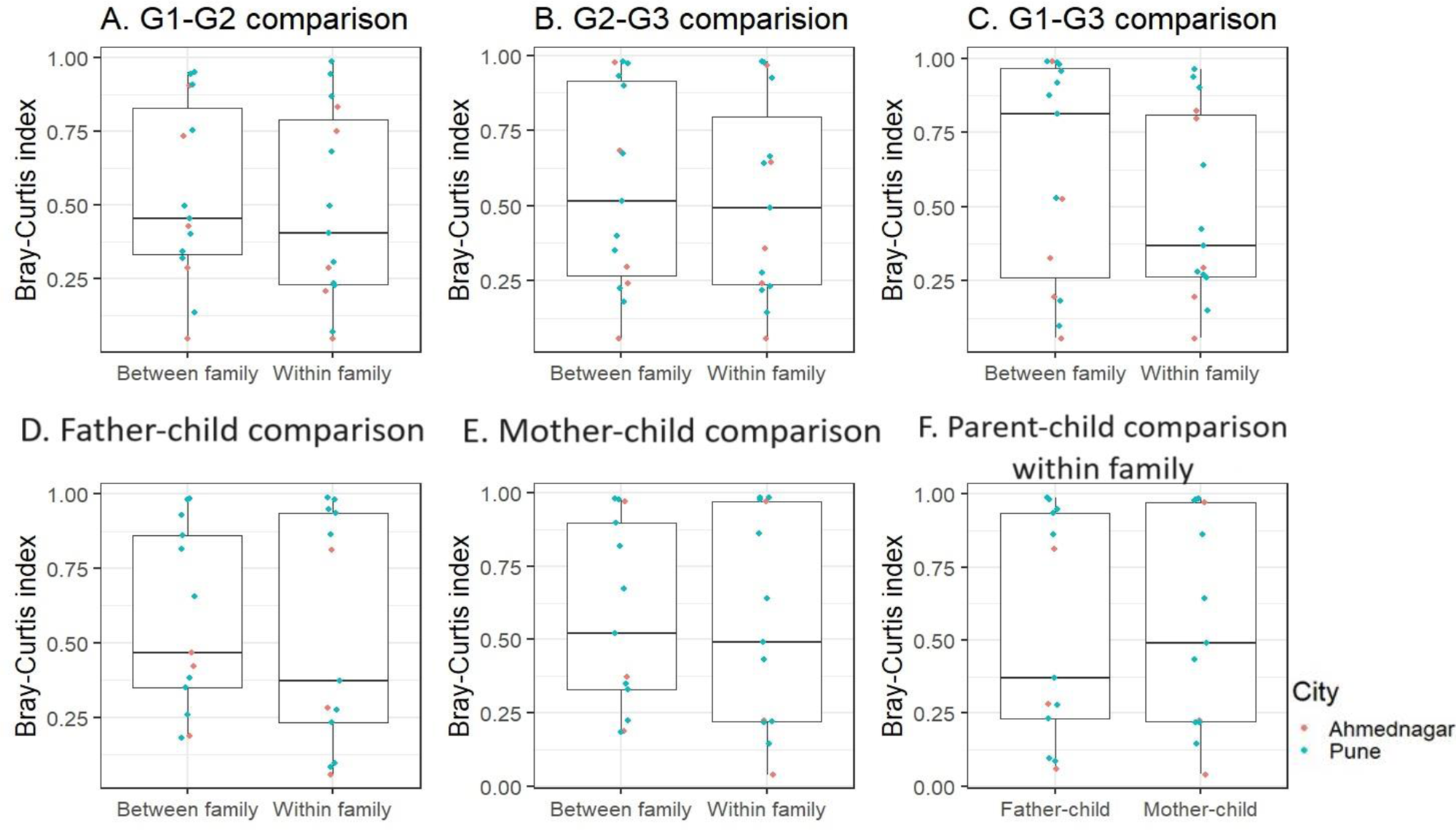
The skin microbiota comparison for within and between family members across three generations A) G1-G2 (grandparent-parent); B) G2-G3 (parent-children); C) G1-G3 (grandparent-child). Separate parent-child comparison (G2-G3) for within and between family D) father-child; E) mother-child; F) parent-child comparison within family. This is compared to the baseline dissimilarity in random adult-child pairs drawn from different families. The differences were not significant (FDR<0.1, Kruskal-Wallis test) for all comparisons.

## Discussion

The composition of skin microbiota is diverse and is influenced by multiple factors. Several studies have indicated predominance of top four phyla-*Actinobacteria*, *Proteobacteria*, *Firmicutes*, and *Bacteroidetes* on healthy human skin (Grice et al., 2009, Schoch et al., 2019, Pammi et al., 2017). Likewise, in the present study, we report the prevalence of phyla *Firmicutes* followed by *Proteobacteria* and *Actinobacteria* on the skin of family members. Our recent results are consistent with earlier studies on the skin microbiota of related individuals from Indian families, which also has revealed an abundance of *Firmicutes, Proteobacteria,* and *Actinobacteria* (Chaudhari et al., 2020). Similarly, our present results align with our previous findings on the skin microbiota of genetically unrelated individuals, wherein we reported predominance of the same phyla-*Firmicutes, Proteobacteria,* and *Actinobacteria* (Potbhare et al., 2022). In contrast, the skin microbiota of infants has been reported to be dominated primarily by *Firmicutes*, followed by abundance of *Actinobacteria,* and *Proteobacteria* (Roux et al., 2022).

Further, analysis of core taxonomic composition revealed the prevalence of commonly colonized skin genera like *Staphylococcus*, *Bacillus*, *Pseudomonas, Anaerococcus*, and *Corynebacterium* in the study population. However, relative abundances of core genera varied from previously reported studies of Roux et al. (2022), where predominance of *Streptococcus* (52.8%) followed by *Cutibacterium* (11.8%), and *Staphylococcus* (8.1%) in infant skin microbiota has been suggested (Roux el al., 2022). Likewise, various studies demonstrated heterogeneous distribution and dominance of *Staphylococcus, Corynebacterium, Streptococcus, and Cutibacterium* genera on skin sites (Saville et al., 2022, Zeeuwen et al., 2012, Ederveen et al., 2020, Byrd et al., 2018). Furthermore, studies based on skin topography and physiology investigated that *Staphylococcus* is dominant in the sebaceous and the moist regions, while *Propionibacterium* dominates only in the sebaceous and *Corynebacterium* only in the moist areas (Catinean et al., 2019). However, the study by Samaras and Hoptroff (2020) reported that irrespective of skin type and site, *Cutibacterium*, *Staphylococcus*, and *Corynebacterium* are present and account for about 45 to 80% of the total skin microbiota (Samaras and Hoptroff, 2020).

### Skin microbiota association with diet

Nutrition is one of the most crucial parameters in modulating skin health and condition. Several studies have been conducted on the gut to understand the influence of diet on microbiota (Bibbo et al., 2016, Schoeler et al., 2019); however, no direct link has been established between skin microbiota composition and diet. We observed moderate association in alpha diversity of vegetarian and mixed (consuming both the plant and animal-based food items, such as eggs, meat, seafood, poultry) diets; however, beta diversity did not show significant differences between the two diet groups. The moderate association in alpha diversity could be due to dietary habits and choices made by the family members. A typical Indian vegetarian diet includes rice, bread, and vegetables as a main course, along with curry made from grains, pulses, sprouts, and dairy products like curd, milk, or cheese. These food choices lead to variations in the amount of fat and fibre consumption and may also be reflected on skin microbiota composition.

Currently, there is growing interest in understanding the connections between the gut and skin microbiota. Mahmud et al. (2022) conducted a study to evaluate the effects of dietary products and probiotics on the microbiota and their interactions between the gut and skin microbiota. They also discussed how antibiotics, prebiotics, medications, long-term diets, and other factors can cause inflammation in gut bacterial communities, which in turn directly affects the skin microbiota. Holscher et al. (2017) also investigated the relationship between gut and skin microbiota and found that dietary choices can cause dysbiosis and lead to various skin diseases. Furthermore, recent research on the facial microbiota of healthy women has suggested that a fasting-mimicking diet can significantly improve skin hydration levels and texture. In addition, the study evaluated how diet affects self-esteem and mental well-being (Maloh et al., 2023).

### Skin microbiota association with age, and sex

Microbiota starts colonizing at the time of birth and changes at early stages, and stabilizes at later stages. A recent microbiota study by Rapin et al. (2023) reported that the delivery mode substantially impacts the colonization of early-life skin microbiota. Further, they investigated that age altered skin microbiota in the first year of life (Rapin et al., 2023). An independent study on the forehead microbiota of healthy Western European women showed significant differences in alpha diversity between elderly group and younger individuals (Jugé et al., 2018). Another study on the facial microbiota of Thai and US age-match males discovered significant differences in middle and elderly age groups. They further investigated that taxa from *Proteobacteria* were more dominant in Thai males, whereas Actinobacteria was predominant in US males (Wilantho et al., 2017). Depending on physiological changes on skin at different ages, pre-pubescent children have high abundances of *Streptococcaceae spp*, *Betaproteobacteria*, and *Gammaproteobacteria* whereas, *Propionibacterium spp.* and *Corynebacterium spp.* were abundant on the skin of post-pubescent individuals (Oh et al., 2012). However, in our present study, we found no significant age-related differences in microbiota diversity (alpha or beta) among individuals of three generations across families. Our results align with the study of Chaudhari et al. (2020), where three generations of Indian patrilineal families were studied to investigate the association of age on skin microbiota. Their comparative analysis revealed no significant differences in alpha diversity of the skin and gut microbiota with three age groups ranging from 3 to >50 years (Chaudhari et al., 2020). Because both of these studies utilize genetically related individuals from Indian families spanning different age ranges, the skin microbiota differences among the three generations’ age groups could be ascribed to either co-habitation or genetic relatedness.

Additionally, gender-specific differences in skin bacteria depend on numerous factors such as skin topography and physiology, e.g., skin thickness, pH, number of hairs, distribution of sweat glands, use of cosmetics, personal hygiene, hormone levels, etc. (Skowron et al., 2021). Earlier, Oh et al. (2012) demonstrated no gender-specific significant association on skin microbiota of the volar forearm (Oh et al., 2012). We did not observe -sex-specific differences in skin microbiota. Ying et al. (2015) demonstrated significant alpha diversity differences in elderly males and females on different skin sites, but microbiota richness was similar between male and female individuals (Ying et al., 2015). On the other hand, axilla studies by Callewaert et al. (2013) showed the predominance of *Corynebacterium* in males and *Staphylococcus* in females (Callewaert et al., 2013). Shami et al., (2019) after analysing cultured bacteria, observed no significant effect of gender on the number of bacteria isolated from four different groups; young people, older adults, males, and females (Shami et al., 2019).

### Skin microbiota association with geographical location

The studies have been conducted globally to explore skin bacterial variations across different geographies by understanding the effects of climatic and environmental conditions leading to spatial patterns. The spatial abiotic factors or climatic conditions are directly related to the host and it affects skin taxonomic composition. This heterogeneity in the microbiota composition can give also spatial information (Ruuskanen et al., 2021). A moderate association with alpha diversity and a borderline significant difference was observed in the beta diversity of the skin microbiota of individuals living in two different geographical locations. These significant differences could be due to slightly varying climatic conditions and urbanization in cities. Geographically Pune covers an area of about 1,110 km^2^ and has a dry climate with an average temperature ranging from 20 to 28°C. Ahmednagar has a hot and semi-arid climate with an average temperature of 15 to 37°C and receives lower rainfall (∼46.3 mm) than Pune (∼67.9 mm). A study by McCall et al. (2020) explained the effect of urbanization on skin microbial compositions by evaluating house structures, outdoor exposure, number of inhabitants of different villages in Amazone (McCall et al., 2020). Urbanization represents a modern lifestyle, good amenities, increased exposure to chemicals, pollution, hygiene, personal care routines, etc. Pune is a metropolitan city, highly urbanized by modern lifestyle, educational development, technology, industries, human resources, health facilities, and transportation system and other differences, compared to Ahmednagar, where the level of urbanization is low as in Ahmednagar, people engaged in agricultural fields, farming, animal husbandry practices Hence they are more exposed to the environment, which might cause differences in bacterial communities. Ying et al. (2015) demonstrated that skin bacteria are significantly influenced by urban and rural populations and residence differences (Ying et al., 2015).

### Skin microbiota variations in family members

Earlier microbiota study of Ross et al. (2017) reported, cohabiting couples shared more similar foot skin microbiota. They further demonstrated that cohabiting partners can be identified based on skin microbiota profile (Ross et al., 2017). Song et al. (2013) compared 60 families with 36 dogs and found that co-habitation resulted in skin microbiota similarities for all comparisons including human-human, dog-dog and human-dog. They further demonstrated that, members of same family shared similar skin microbiota than members of different households, stating shared environment may homogenize skin microbiota through contact with common surfaces (Song et al., 2013). In our present study, we have observed significant variations in skin microbiota between different families, while significant differences within members of the same family were not observed. This variation could be attributed to cohabitation or the genetic relatedness among family members spanning three generations. Consequently, it raises the question of whether skin microbiota carries a distinctive familial signature. However, drawing such a conclusion would necessitate a dedicated study with a larger sample size and a greater number of individuals within the same generation. Similar to what has been suggested previously for gut microbiota, genetically related individuals might exhibit more similarity in their gut microbiome compared to genetically unrelated individuals, regardless of co-habitation patterns (Turnbaugh et al., 2009). Yatsunenko et al. (2012) observed that teenagers share more similar faecal microbiota with their parents than with unrelated adults (Yatsunenko et al., 2012). Study by Si et al. (2015) on the skin microbiota diversity of monozygotic and dizygotic twins suggested that variation is due to host genetics, age, and skin pigmentations (Si et al., 2015). A recent study on facial skin microbiota investigated genetic factors that influence aging-related pathways causing skin microbial differences in healthy Italian women in three age groups (younger, middle-aged, and older) (Russo et al., 2023). However, an in-depth analysis of heritability of a skin microbiota across generations will require larger number of families and samples per generation.

### Study limitations

The objective of the current study was to understand the differences and similarities in skin microbiota of family members spanning three generations. However, one of the major limitations of the present study is that a majority of the families included had either only one grandparent or a single sibling. Consequently, there was variability in the number of family members present across the three generations within the enrolled families. The outcomes of this study could have been more robust had the analysis encompassed more than two individuals per generation (such as cousins) in order to encompass the full spectrum of genetically related and unrelated family members. Moreover, our dataset is based on only 15 families, which has limited statistical power to detect significant differences. The inclusion of a greater number of families is imperative to more comprehensively explore the intricate interplay between the similarity of skin microbiota, genetic relatedness, and the diverse confounders.

## Conclusion

The present study highlighted the composition of skin microbiota from Indian families which was found to be typically characterized by the presence of phyla such as *Firmicutes*, *Proteobacteria*, and *Actinobacteria*, and genera *Staphylococcus*, *Bacillus*, *Pseudomonas, Anaerococcus*, and *Corynebacterium*. We observed moderate alpha diversity association for diet and geographical location, and borderline significant beta diversity with geographical location on skin microbiota among the participating families. No significant differences were observed when within and between family comparisons were made across generations and within family skin microbiota composition was more similar/less diverse than members from different families. For a comprehensive understanding of the interplay between genetics and cohabitation, it is imperative to undertake future research with larger number of families having a greater number of siblings spanning multiple generations.

## Authors’ contribution

RA conceived the study; RA, LL, EM, AR, and RP involved in the study design discussion; RP performed the experiments, methodology, statistical analysis, data analysis and wrote first draft of the manuscript; LL coordinated the statistical analysis and data science; RA, LL supervised study and supported writing first draft of the manuscript; LL, RA, EM and AR reviewed and edited the manuscript. All the authors read and approved the final manuscript.

## Acknowledgments

Authors are grateful to the volunteers for their participation in this study. The authors thank MHRD-SPARC (Government of India) for financial support and Department of Zoology, Savitribai Phule Pune University, for providing infrastructure and equipment facilities. Authors also thank Rashtriya Uchchtar Shiksha Abhiyan (RUSA, Savitribai Phule Pune University) for student fellowship.

## Code availability

The source code for the analysis is available on GitHub and accessed via permanent DOI https://doi.org/10.5281/zenodo.10297062

## Data availability

All raw sequences data were deposited and available at European Nucleotide Archive with Project accession no-PRJEB44216 from ERS6234946 to ERS6234956, ERS6234959 to ERS6234976, ERS6234984 to ERS6234989 and Project accession no-PRJEB62887 from ERS15575930 to ERS15584034

