## Supplementary figure 1 for "Skin microbiota variation in Indian families"

*First Author

Renuka Potbhare, Ph.D student, Department of Zoology, Savitribai Phule Pune University, Pune – 411 007. Maharashtra, India.

#Corresponding Authors

1. Leo Lahti, Professor, Department of Computing, University of Turku, Finland.

2. Richa Ashma, Associate Professor, Department of Zoology, Savitribai Phule Pune University, Pune – 411 007. Maharashtra, India.

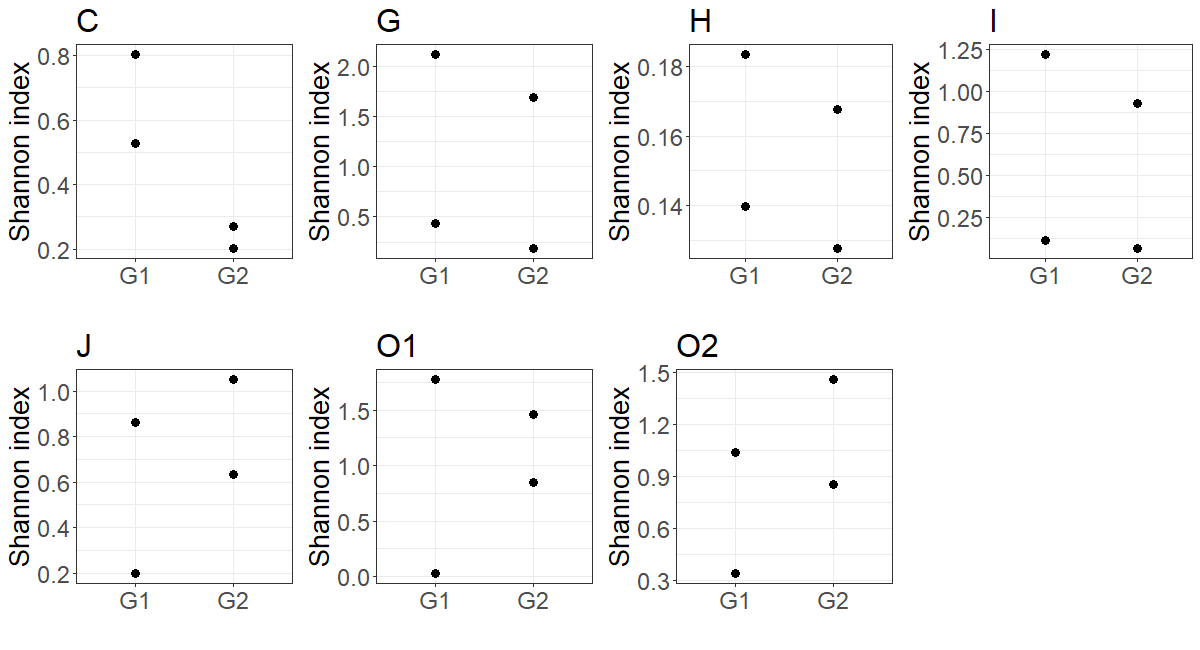

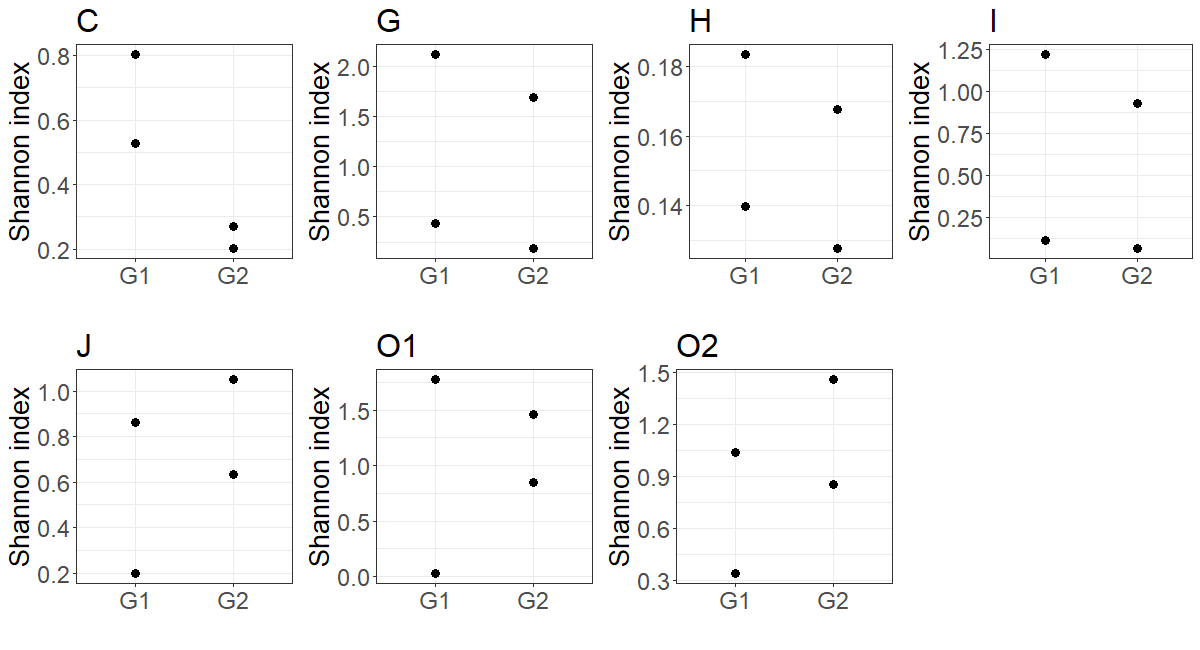

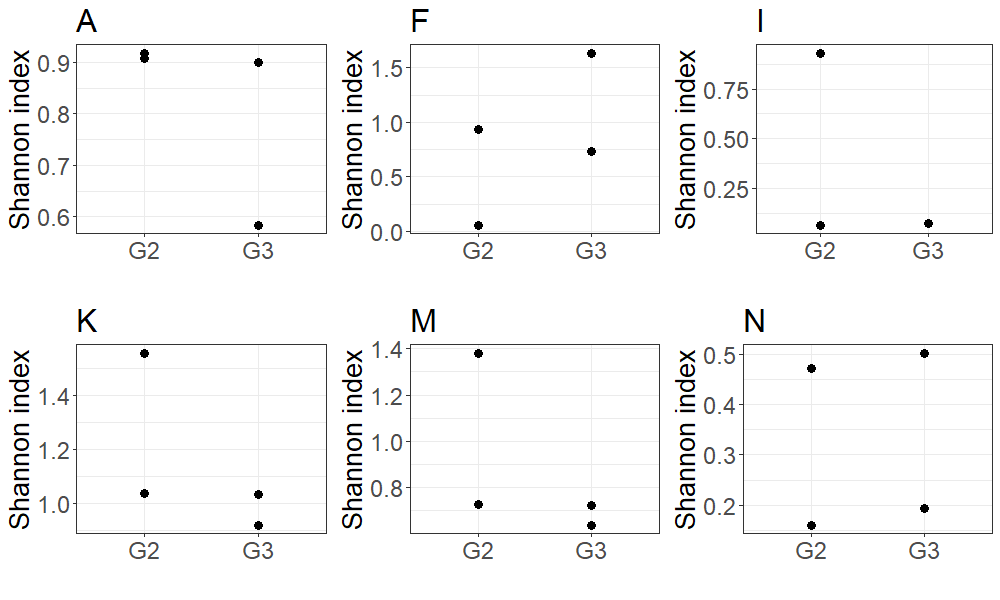

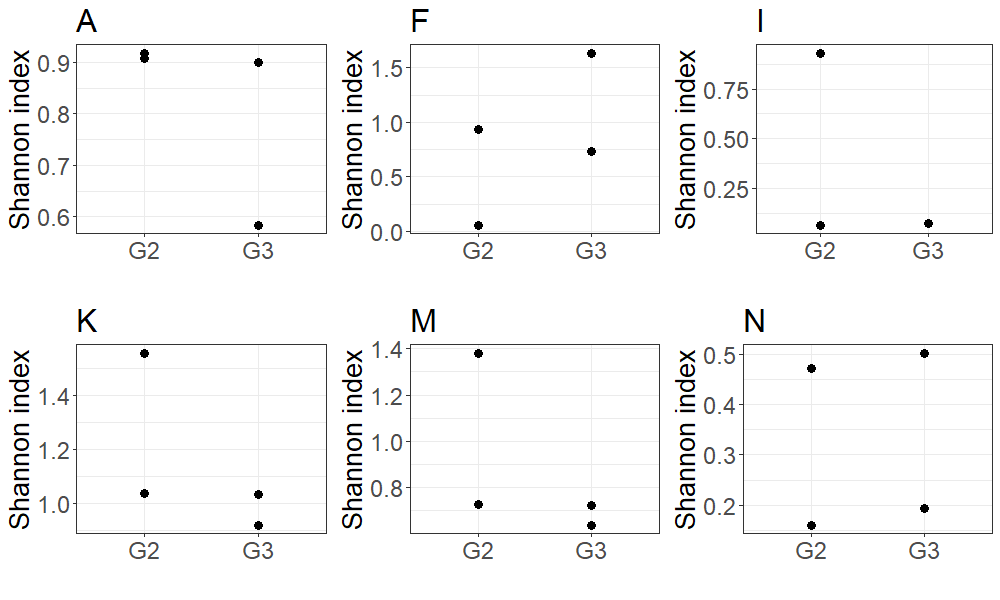

Supplementary figure 1: Within family inter-generational differences. First panel represents G1-G2 intra-family comparison and second panel represents G2-G3 intra-family comparison. The analysis is carried out only on families having both the members per generation hence the comparison differs among the generations across families. p-values were adjusted for multiple testing (Kruskal-Wallis test, FDR<0.1) for all the comparisons.
